# Cellular-resolution OCT reveals layer-specific retinal mosaics and ganglion cell degeneration in mouse retina in vivo

**DOI:** 10.1101/2025.08.01.668001

**Authors:** Tae-Hoon Kim, Robby Weimer, Justin Elstrott

**Author notes:** **Correspondence**: Justin Elstrott, Department of Translational Imaging, Genentech, Inc., 1 DNA Way, South San Francisco, CA 94080, USA.

## Abstract

Cellular-resolution retinal imaging in preclinical mouse models is limited by optical aberrations and speckle noise that prevent visualization of individual cells. We developed a wavefront sensorless adaptive optics optical coherence tomography (WSAO-OCT) platform that addresses these challenges by combining real-time aberration correction with multi-volume averaging. This integrated approach enhanced image contrast by 61% and sharpness by 55%, while averaging 50 volumes effectively reduced speckle noise. Our system enabled visualization of individual cells across all major retinal layers and the retinal pigment epithelium in living mice. Immunohistochemical validation within the retinal ganglion cell (RGC) layer confirmed that 95% of optically detected cells corresponded to actual RGCs, confirming cell-type specificity. We demonstrated the method’s translational potential by tracking RGC loss following optic nerve injury. The platform detected significant cell degeneration within three days post-injury, earlier than traditional thickness-based methods, quantifying a decline from 4,407 to 1,683 cells/mm^2^ (62% loss) over seven days. These results establish WSAO-OCT as an effective, non-invasive tool for cellular-level investigation of retinal pathologies, suitable for longitudinal monitoring and therapeutic evaluations in preclinical models.

## Introduction

Recent advances in adaptive optics optical coherence tomography (AO-OCT) have demonstrated cellular-resolution imaging in human retina^1^. Cellular resolution imaging enhances our understanding of retinal pathophysiology by enabling direct observation of early pathological changes, which may facilitate a shift in diagnostic and therapeutic monitoring from tissue to cellular resolution. However, translation of these capabilities to preclinical models presents significant technical challenges. This challenge is particularly acute in mouse models, which, despite their widespread use in retinal research, pose unique optical and physiological barriers to achieving cellular-resolution imaging in vivo^2,3^. These barriers persist even though modern OCT systems achieve theoretical spatial resolutions sufficient to resolve individual cells such as mouse retinal ganglion cell (RGC) somata (5-10 µm diameter).

The practical implementation of cellular-resolution OCT in mouse retina faces two fundamental optical barriers beyond the challenge of low refractive index contrast between adjacent cells. First, ocular aberrations introduce significant wavefront distortions that degrade image quality. Although the mouse eye’s high numerical aperture should theoretically confer high resolving power, this optical advantage is paradoxically compromised by severe aberrations that scale with numerical aperture, resulting in image degradation rather than enhanced resolution^4,5^. Second, the coherent detection mechanism intrinsic to OCT generates speckle noise, stochastic interference patterns. This speckle pattern effectively masks the underlying cellular structures, as the grain size is comparable to cellular features^6,7^. The convergence of these optical phenomena, aberration-induced blur and coherent speckle noise, fundamentally limits contrast discrimination in densely packed nuclear layers.

To overcome these limitations, we devised an integrated imaging platform that merges three synergistic components: (i) a custom high-resolution spectral-domain OCT (SD-OCT), (ii) a wavefront-sensorless adaptive-optics (WSAO) module that corrects ocular aberrations, and (iii) a hierarchical multi-volume averaging algorithm that suppresses speckle noise. This platform resolves single cells throughout the living mouse retina, from ganglion cells to photoreceptors and the retinal pigment epithelium (RPE). We demonstrate its translational utility by longitudinally tracking RGC soma degeneration after optic nerve crush (ONC) injury, detecting a ∼23% loss within three days, thereby establishing WSAO-OCT as a sensitive tool for pre-clinical studies of retinal disease and therapeutic evaluation.

## Results

### WSAO Effect on OCT Image Quality

We first evaluated the effectiveness of WSAO in correcting ocular aberrations in mouse eyes by assessing OCT image quality improvements across retinal quadrants. The field of view (FOV) was centered on the optic nerve head (ONH), with small patches (310 × 310 μm) selected from four quadrants (inferior-nasal, inferior-temporal, superior-nasal, superior-temporal) (Fig. 1a, b). En face OCT images were acquired from 39 regions of interest (ROI) across 8 mice, with a variable number of ROIs per mouse, both without AO correction (flattened deformable mirror) and with one round of Zernike modal optimization. WSAO significantly enhanced image quality across all retinal quadrants, with contrast improvements of 64%, 60%, 40%, and 67% in the inferior-nasal, inferior-temporal, superior-nasal, and superior-temporal quadrants, respectively (improvement ratios of 1.64 ± 0.30, 1.60 ± 0.21, 1.40 ± 0.38, and 1.67 ± 0.31) (Fig. 1c1). Similarly, sharpness improved by 60%, 55%, 38%, and 59% in the inferior-nasal, inferior-temporal, superior-nasal, and superior-temporal quadrants, respectively (improvement ratios: 1.60 ± 0.28, 1.55 ± 0.20, 1.38 ± 0.37, and 1.59 ± 0.28) (Fig. 1c2). Overall, WSAO-based aberration correction yielded 61% enhancement in contrast (ratio: 1.61 ± 0.31) and a 55% increase in sharpness (ratio: 1.55 ± 0.29), demonstrating effective compensation for optical aberrations in the mouse eye and enabling improved visualization of retinal structures (Fig. 1d).

**Figure 1.**
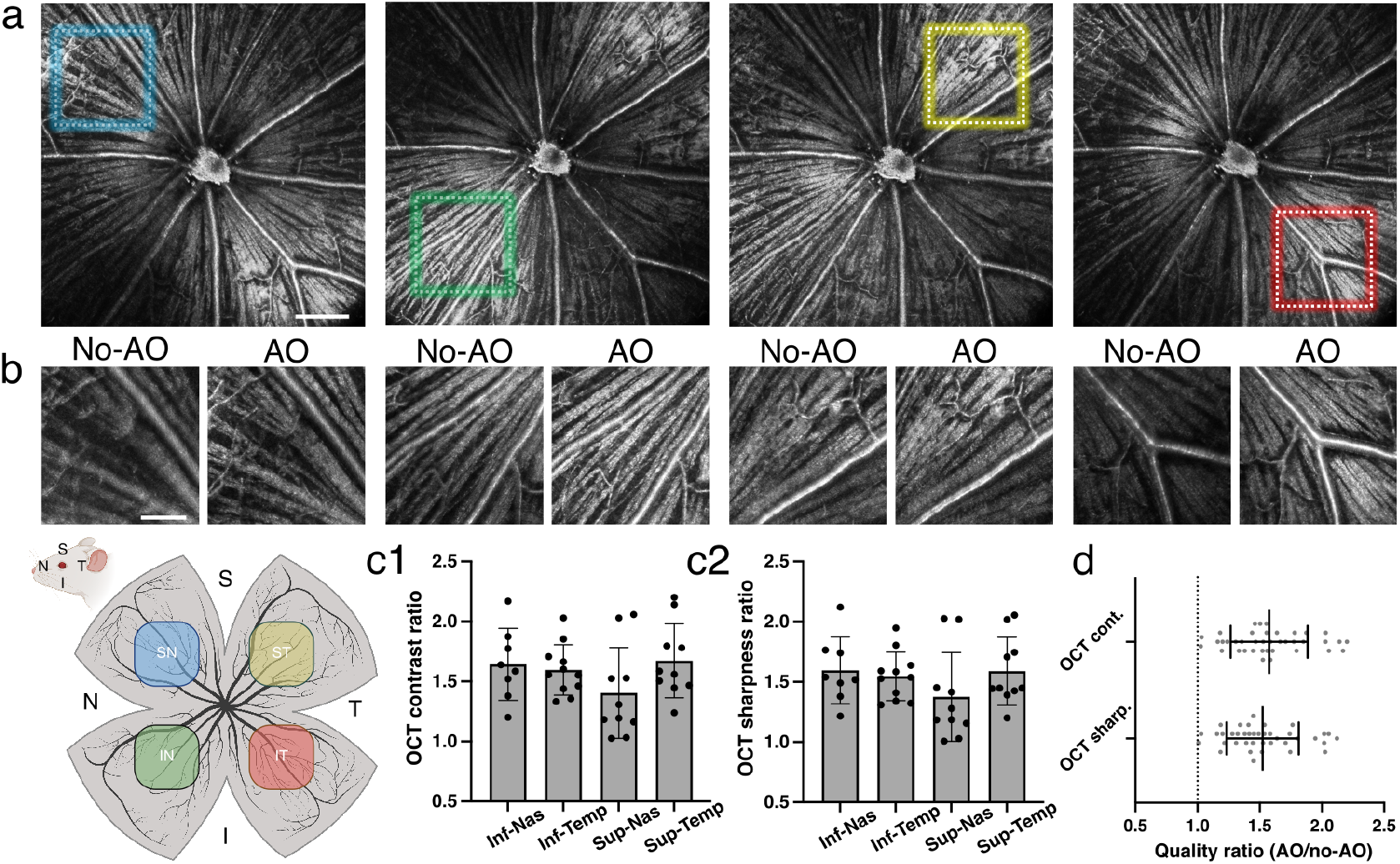
WSAO enhances OCT image quality across the mouse retina. (a) Representative en face images of the retinal nerve fiber layer around the ONH, with colored boxes indicating the small patches selected for WSAO optimization. (b) Comparative patch images before and after WSAO optimization demonstrating a clear improvement in image contrast and feature visibility. (c1, c2) Key image quality metrics, such as contrast ratio and sharpness ratio, plotted for four retinal quadrants to assess spatial uniformity of the correction. (d) Global average of these quality metrics, quantified from a total of 39 ROIs from 8 mice. Scale bars, 200 µm in (a); 100 µm in (b).

### Multi-Volume Averaging Effect on OCT Image Quality

We investigated speckle noise reduction through volume averaging, focusing on cellular visibility in the RGC layer. Data were collected from a patch of 310 × 310 μm in the retinal nerve fiber layer (RNFL), with 200 volumes acquired sequentially after aberration correction. Representative *en face* images (Fig. 2a) demonstrated that single volumes exhibited prominent speckle noise obscuring cellular details, while averaging 5 volumes significantly reduced noise and averaging 20 volumes revealed the cellular mosaic. Averaging 50 volumes further enhanced visibility with minimal noise, though improvements beyond 50 volumes were negligible. Quantitative analysis of normalized speckle contrast (Fig. 2b) confirmed rapid noise reduction from 1.00 (single volume) to 0.60 ± 0.08 (5 volumes) and 0.49 ± 0.11 (20 volumes), plateauing at 0.46 ± 0.10 (50 volumes). Structural similarity index (SSIM) analysis (Fig. 2c), using the 200-volume average as the ground truth reference, showed progressive improvement from 0.13 ± 0.04 (single volume) to 0.46 ± 0.10 (5 volumes), 0.79 ± 0.08 (20 volumes), and 0.85 ± 0.06 (50 volumes), plateauing near 0.93 ± 0.04 (75 volumes). Line intensity profiles across three cell bodies (Fig. 2d1) demonstrated high similarity among 50-, 100-, and 200-volume averages. Cross-correlation analysis among these line intensity profiles (Fig. 2d2) showed coefficients increasing from 0.33 ± 0.09 (single volume) to 0.58 ± 0.11 (5 volumes), 0.81 ± 0.08 (20 volumes), and reaching 0.94 ± 0.03 at 75 volumes, confirming plateau between 50-75 volumes. These results indicate that most image quality improvements occur within 50 volumes, with metrics plateauing between 50-75 volumes. This range represents the optimal balance between image quality and acquisition efficiency for visualizing cellular architecture in OCT imaging.

**Figure 2.**
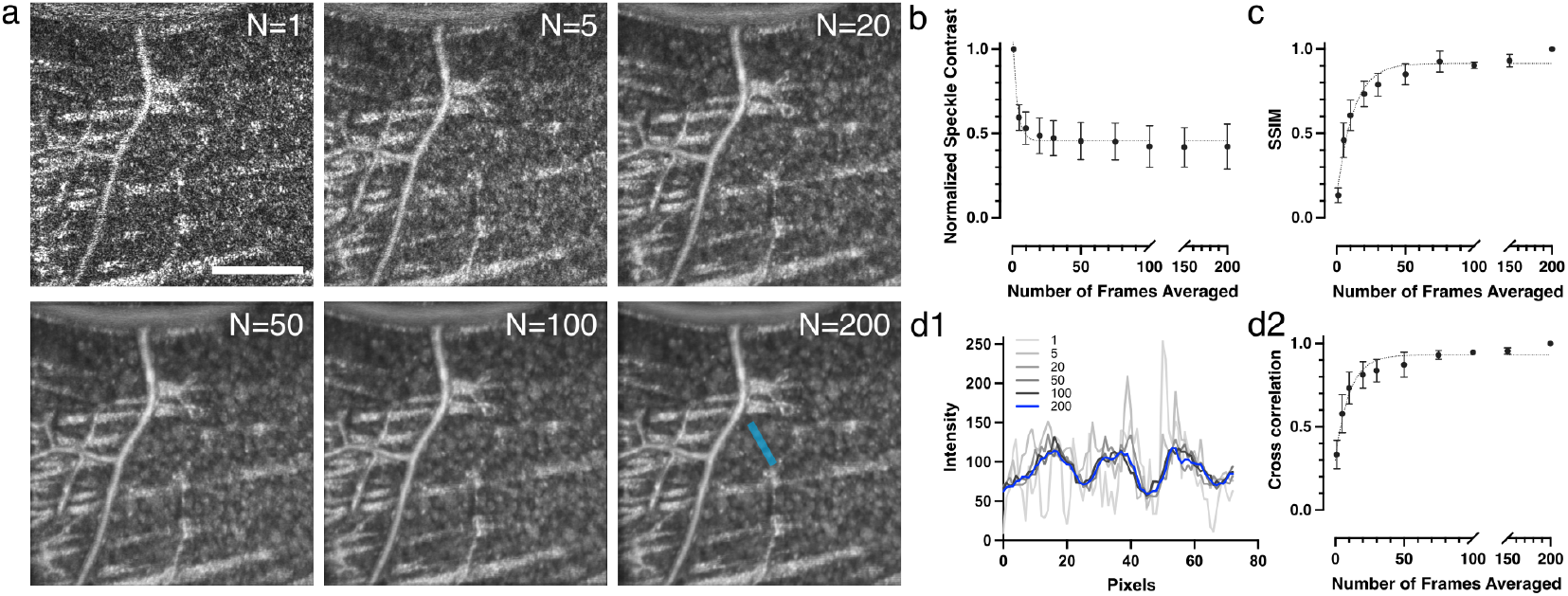
Multi-volume averaging enhances OCT image quality and cellular visibility in the RGC layer. (a) Representative *en face* images from the same retinal location, demonstrating the progressive reduction of speckle noise and improved visibility of the cellular mosaic as the number of averaged volumes increases. (b) Quantitative analysis of normalized speckle contrast plotted against the number of averaged volumes, showing a rapid initial decrease that subsequently plateaus. (c) Structural Similarity Index analysis, using the 200-volume average as a reference, illustrates the improvement in image fidelity with increased averaging. (d) Effect of volume averaging on the quality of cellular line intensity profiles. (d1) Line intensity profiles across three representative cell bodies, generated from *en face* images created by averaging 1, 5, 20, 50, 100, and 200 volumes. The profile location is exemplified by the blue line in panel (a) at N=200 image. (d2) Cross-correlation coefficients of these profiles relative to the 200-volume average, confirming that structural details stabilize after approximately 75 volumes. Data in plots are presented as mean ± SD. Sample sizes were N = 18 ROIs from 7 mice for (b), N = 9 ROIs from 3 mice for (c), and N = 10 ROIs from 3 mice for (d2). Scale bar, 100 µm.

### WSAO OCT Reveals Cellular Mosaics Across Retinal Layers

We applied volume averaging with WSAO to visualize cellular mosaics in all retinal nuclear layers (RGC, inner nuclear layer (INL), outer nuclear layer (ONL)) and the RPE. A tunable-focus lens enabled precise focal plane positioning, with WSAO implemented at each layer. Representative B-scan images (Fig. 3a) demonstrate focal plane adjustments through dioptric shifts, with corresponding retinal intensity profiles (Fig. 3b). Relative to the RGC layer baseline, dioptric shifts of 2.4D, 3.2D, and 4.0D were required to focus on the INL, ONL, and RPE layers, respectively. Average intensity *en face* images generated from the registered and averaged volumes at four separate focal depths (100 volumes each) revealed layer-specific cellular morphologies (Fig 3c). The RGC layer (Fig. 3c1) displayed densely packed, round somas with uniform reflectance. The INL (Fig. 3c2) exhibited heterogeneous cell populations, including small highly reflective cells and larger dimmer somas, consistent with the diverse cell types in this layer. The ONL (Fig. 3c3) consisted of small, uniformly distributed cell bodies with variable reflectance patterns. The RPE layer of pigmented C57BL/6J mice (Fig. 3c4) exhibited regularly spaced circular hyporeflective regions surrounded by hyperreflective regions. These features likely represent RPE nuclei (hyporeflective) and surrounding melanosomes (hyperreflective), as this mosaic pattern was absent in albino mice (Supplementary Fig. S1). A fly-through visualization of these distinct layers is available as Supplementary Movie S1.

**Figure 3.**
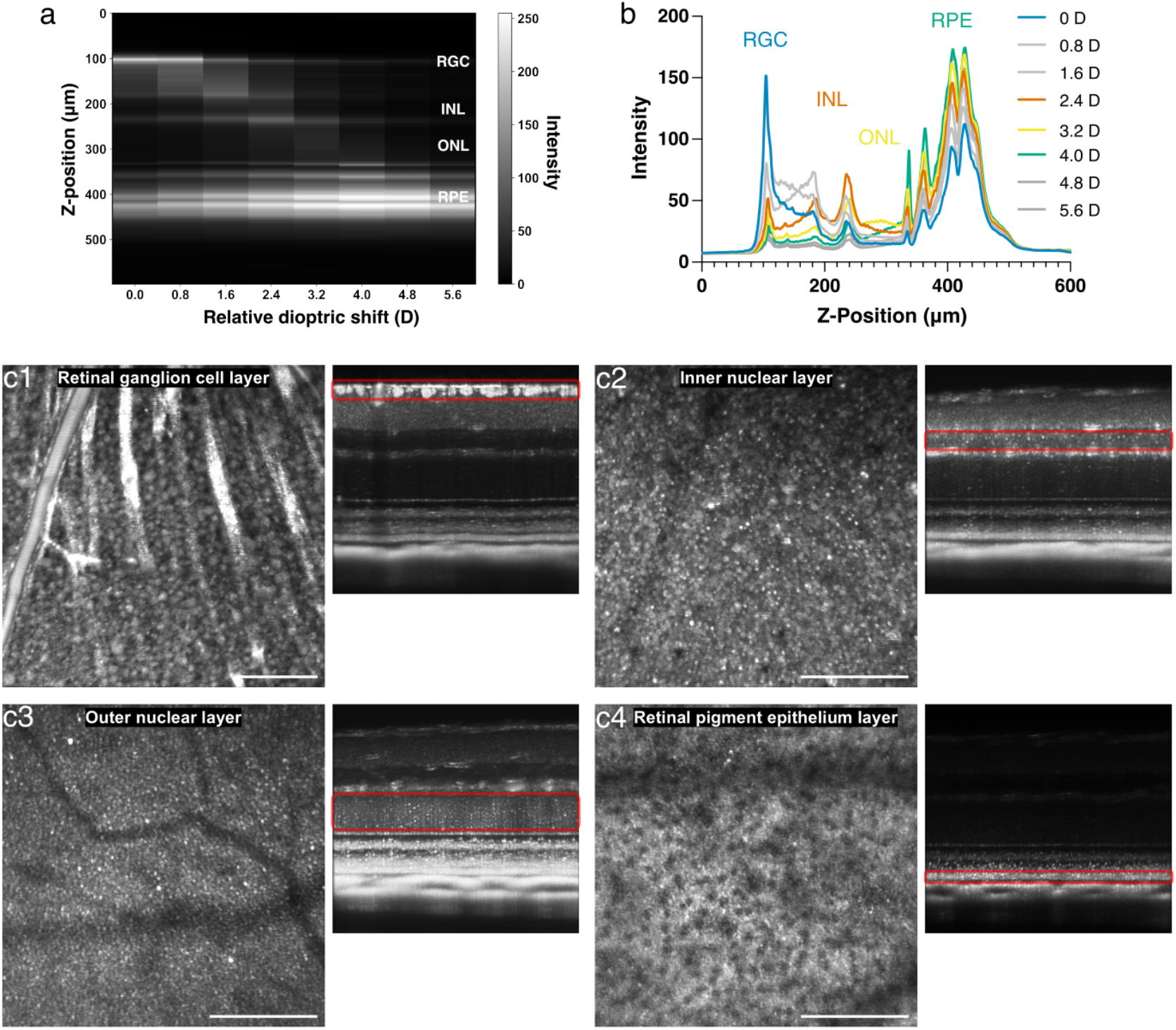
Layer-specific visualization of cellular mosaics across the mouse retina using WSAO-OCT. (a) Representative B-scans demonstrating precise focal plane adjustments to target different retinal layers through applied dioptric shifts. (b) Corresponding axial intensity profiles for each focal plane shown in (a). (c) High-resolution en face images of distinct cellular mosaics revealed after WSAO and multi-volume averaging. Panels show the RGC layer (c1), INL (c2), ONL (c3), and the RPE in a pigmented C57BL/6J mouse (c4). The B-scan next to each *en face* image shows the corresponding focal plane and enhanced contrast for that layer. Scale bars, 100 µm.

### WSAO OCT Detects Early Retinal Ganglion Cell Loss in Optic Nerve Injury Model

To demonstrate the translational capability of our imaging workflow for quantitative cell analysis in disease models, we applied this technique to the ONC model, an established model of acute traumatic injury that induces approximately 50% RGC loss within 7 days^8^. Three mice underwent unilateral ONC in the left eye, with longitudinal imaging performed at baseline and post-injury timepoints. For each retina, OCT data were acquired from two adjacent patches in the superior quadrant, and the resulting cell counts were averaged to provide a single value for quantifying cell counts. Figure 4a displays representative longitudinal *en face* images of the RGC layer, illustrating progressive cell loss post-injury. The quantitative analysis also revealed progressive cell loss following optic nerve injury, with clear evidence of neurodegeneration detectable as early as 3 days post-crush (Fig. 4b). Cell counts in the RGC layer declined progressively across the five matched time points (repeated-measures ANOVA: F(1.48, 2.95)= 127.2, P = 0.0015; ε = 0.369). Cell density decreased from baseline 424 cells to 414 cells at 1 day (P = 0.236), then significantly to 327 cells at 3 days (P = 0.021), 226 cells at 5 days (P = 0.022), and 162 cells at 7 days (P = 0.0028), representing a 62% reduction from baseline. When normalized to area, cell density declined from 4,407 cells/mm^2^ at baseline to 4,310 cells/mm^2^ at 1 day, 3,397 cells/mm^2^ at 3 days, 2,348 cells/mm^2^ at 5 days, and 1,683 cells/mm^2^ at 7 days. Ganglion cell complex (GCC) thickness exhibited a parallel but delayed trajectory (repeated-measures ANOVA: F(1.11, 2.22)= 148.5, P = 0.0045; ε = 0.277) (Fig. 4c). GCC thickness remained stable from baseline 79.1 µm to 80.6 µm at 1 day (P = 0.086) and 79.9 µm at 3 days (P = 0.622), before showing significant reductions to 77.3 µm at 5 days (P = 0.0034) and 72.5 µm at 7 days (P = 0.0055), corresponding to an 8% decrease. Collectively, these data suggest that in vivo cell counting in the RGC layer may be a more sensitive early marker, detecting significant loss by 3 days post-injury while GCC thickness changes only became significant at 5 days, although both metrics converge to show substantial degeneration within the first post-injury week.

**Figure 4.**
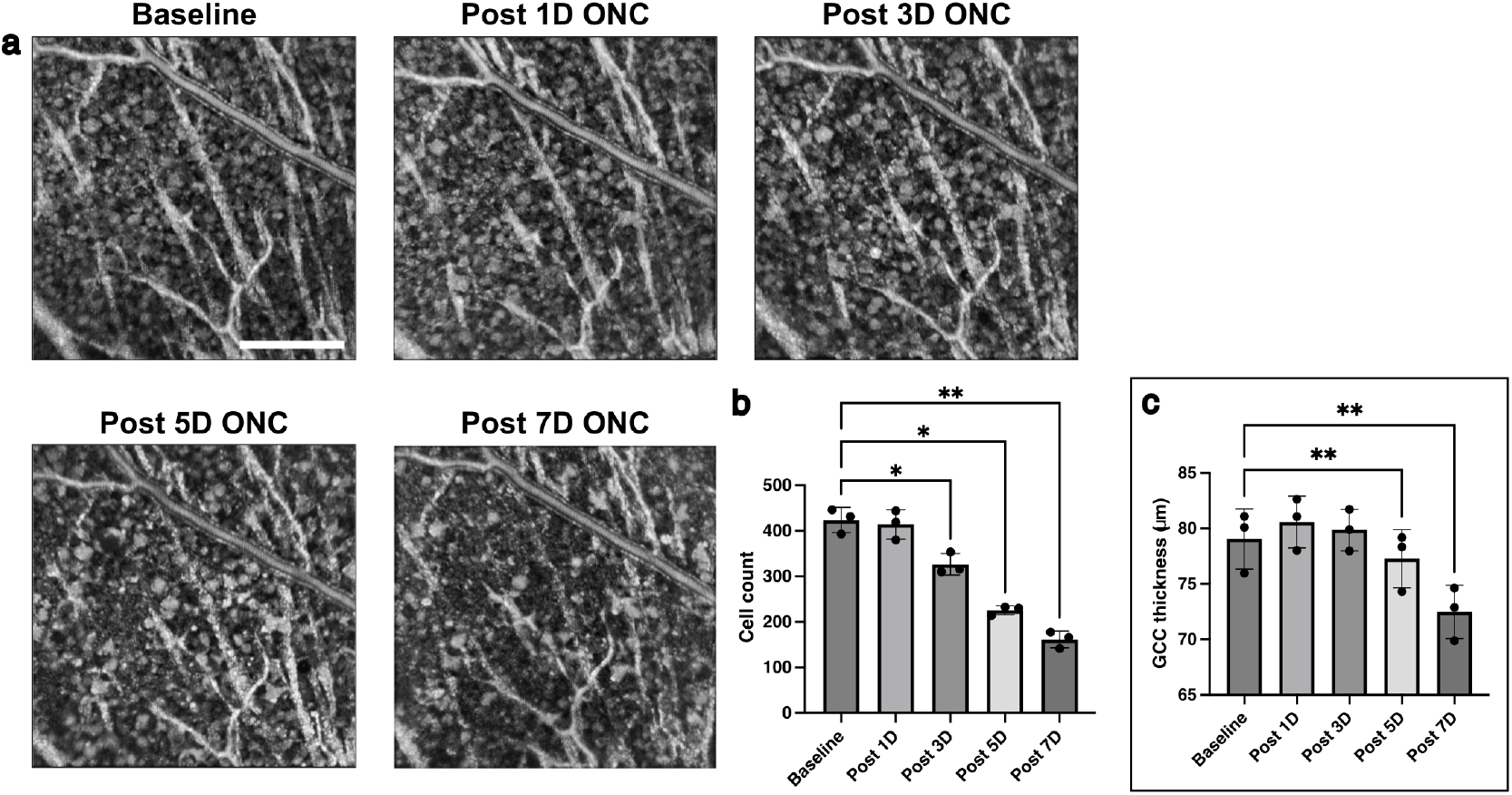
Longitudinal tracking of soma loss and GCC thinning following optic nerve crush. (a) Representative longitudinal *en face* images of the RGC layer from the same retinal location, shown at baseline and at 1, 3, 5, and 7 days post-ONC. The images qualitatively illustrate the progressive loss of cell bodies over the first week following injury. (b) Quantification of cell counts over the 7-day period. The number of visible somas shows a significant decline beginning at 3 days post-injury. (c) Quantification of GCC thickness over the same time course. In contrast to direct cell counting, a significant reduction in GCC thickness is detected later, starting at 5 days post-injury. Data are presented as mean ± SD (N = 3 mice). Asterisks denote statistically significant differences compared to baseline, as determined by post-hoc tests following a repeated-measures ANOVA (*P < 0.05, **P < 0.01). Scale bar, 100 µm.

### WSAO OCT Cellular Signals Correspond to RGC Somas

To validate the cellular identity of high-contrast structures observed in the longitudinal ONC images, we performed immunohistochemical analysis on whole-mount retinas following OCT imaging. Triple staining was performed using NeuroTrace (pan-neuronal marker, Fig. 5a1), RBPMS (RGC-specific marker, Fig. 5a2), and CD31 (vascular marker, Fig. 5a3) on retinas 7 days post-ONC. Retinal vasculature labeled with CD31 served as anatomical landmarks for precise co-registration between OCT and fluorescence images. This co-registration was also used to calibrate the OCT FOV (Supplementary Fig. S2). Spatial correlation analysis revealed strong correspondence between high-contrast cells in OCT image and RBPMS-positive RGC somas (Fig. 5b). Quantitative comparison within matched fields of view (310 × 310 µm) from 4 mice also demonstrated agreement (Fig. 5c): OCT identified 154 ± 18 cells, while RBPMS staining revealed 163 ± 19 cells. In contrast, NeuroTrace labeled 620 ± 83 cells, a number nearly four times higher, capturing the entire neuronal population including displaced amacrine cells. The near-identical counts between OCT and RBPMS (154/163 = 95% detection rate) confirm that WSAO OCT visualizes RGC somas with minimal contribution from non-RGC neurons. Collectively, the spatial correlation analysis and the high concordance of cell counts with the RGC-specific marker RBPMS provide strong evidence that the high-contrast structures in the RGC layer are predominantly RGC somas.

**Figure 5.**
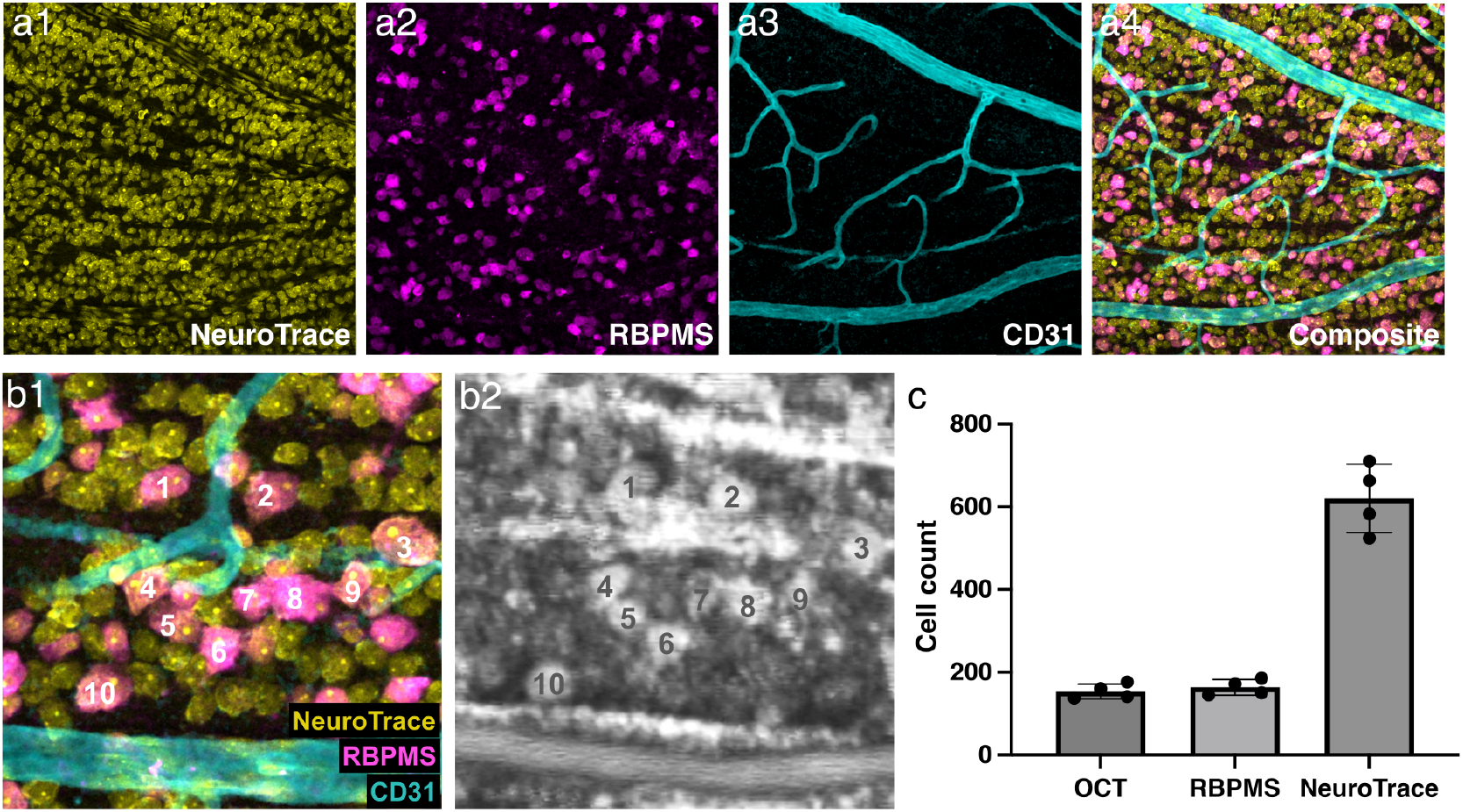
Validation of high-contrast OCT signals as RGC somas using co-registered immunohistochemistry. (a) Representative images of a whole-mount retina after triple-staining for all neurons (NeuroTrace, yellow), RGCs specifically (RBPMS, magenta), and blood vasculature (CD31, cyan), alongside the composite image (a4). (b) Magnified view demonstrating the direct spatial correspondence between the RBPMS-positive cells in the co-registered fluorescence composite (b1) and the high-contrast somas in the in vivo OCT image (b2). Numbered cells highlight the one-to-one anatomical match. (c) Quantitative comparison of cell counts at 7 days post-injury within matched fields of view. The cell density measured by OCT closely matches that of the RGC-specific marker RBPMS, and is substantially lower than the total neuronal count provided by NeuroTrace. Data are presented as mean ± SD from N = 4 mice.

## Discussion

Here we demonstrate that integrating WSAO with volumetric averaging enables visualization of cellular mosaics across all neuronal layers and RPE in living mice. Our systematic analysis revealed complementary benefits: WSAO corrected optical aberrations to enhance image sharpness and contrast across retinal regions, while volume averaging suppressed speckle noise to reveal previously obscured cellular details. Through quantitative metrics including normalized speckle contrast, SSIM, and cross-correlation analysis, we determined that averaging approximately 50 volumes optimally balanced image quality improvement with practical acquisition time constraints. Immunohistochemical validation confirmed that high-contrast structures in the RGC layer corresponded specifically to RGC somas with 95% concordance to RBPMS staining, demonstrating selective visualization despite the mixed neuronal population. Application to an ONC model further validated the technique’s sensitivity for detecting pathological changes, successfully quantifying progressive RGC loss with sufficient precision for longitudinal disease monitoring. These findings establish WSAO OCT as a powerful platform for non-invasive investigation of retinal neurodegeneration at cellular resolution.

Our imaging strategy builds upon two distinct but complementary lines of prior research that have addressed the optical aberrations and speckle noise in the mouse retinal imaging. Jian et al. pioneered Shack-Hartmann-guided AO-OCT, which corrected optical aberrations to achieve near-diffraction-limited imaging (wavefront error ∼50 nm) and substantially enhanced signal intensity^4^. Subsequent sensorless AO implementations by Jian et al.^9^ and Wahl et al.^10^ similarly optimized Zernike modes to enhance OCT intensity and sharpness. More recently, Zhang et al. developed a high-speed AO-SLO-OCT platform that significantly reduced wavefront error and improved resolution^11^. Consistent with these pioneering studies, our WSAO implementation increased image sharpness by 55% and contrast by 61%. A separate line of investigation has focused specifically on suppressing speckle noise through volume averaging to visualize individual cells in the mouse retina. Zhang et al. first demonstrated that averaging up to 150 OCT volumes could reveal RGC somas, establishing that speckle contrast reduction plateaued after ∼50 volumes^12^, a finding consistent with our own analysis. Their follow-up work further validated this temporally-averaged OCT approach for label-free cellular imaging through co-localization with fluorescent reporters^13^. While prior work established the feasibility of cellular visualization through averaging, its lack of AO meant that inherent resolution and contrast were limited by the system’s uncorrected optics. Our study directly bridges this gap by synergistically combining both strategies. By integrating adaptive optics with volume averaging, we achieve superior cellular visualization that could enable more precise longitudinal tracking and pathological assessment in mouse models of neurodegeneration.

This combined approach builds on principles pioneered in human imaging studies, highlighting the translational relevance of our findings. Liu et al. visualized RPE mosaics in human subjects by averaging 24-35 AO-OCT volumes, revealing distinct cellular features including hyporeflective nuclei^14^, findings that parallel our observations in pigmented mice. More relevant to our RGC findings, Liu et al. employed high-speed AO-OCT to image ganglion cell mosaics across the human retina^15^. By averaging 100-160 registered volumes, they achieved significant contrast improvement that revealed over 42,000 individual RGC somas without exogenous labeling. Their optimization studies strongly align with our own: fewer than 50 volumes produced inadequate contrast, while ∼100 volumes provided an optimal balance, consistent with our findings. The consistency of imaging parameters and results across species validates the technical performance of our WSAO-OCT platform, establishing it as a valuable tool for bridging preclinical mouse studies and human clinical applications.

A key finding of our study is the selective visualization of a specific cellular subpopulation within the RGC layer. The high-contrast somas we identified not only showed 95% concordance with the RGC-specific marker RBPMS, but their quantified density provides further evidence for their identity. The mouse RGC layer contains RGCs (41% of total neurons) and displaced amacrine cells^16^. Our measured baseline density of 4,407 cells/mm^2^ for the high-contrast somas is consistent with the known density of RGCs. Foundational census studies by Jeon et al.^16^ and Salinas-Navarro et al.^17^ report a whole-retina average RGC density of approximately 3,300 cells/mm^2^ in C57BL/6J mice. These studies also describe a non-uniform distribution, with a temporal-superior “visual streak” where local RGC densities can peak at over 8,500 cells/mm^2^. Therefore, our measurement of ∼4,400 cells/mm^2^ is consistent with imaging a retinal region of intermediate-to-high RGC density.

Further evidence for this identification as RGCs comes from our longitudinal ONC experiment. Following the ONC injury, which selectively induces RGC apoptosis, the density of these high-contrast somas in OCT decreased to 1,683 cells/mm^2^. This post-injury density shows close agreement with recent findings by Zapadka et al., who reported a similar density of 1,700 ± 321 RBPMS-positive cells/mm^2^ at 7 days post-crush^18^. This specific loss of the visualized cells in a model of RGC degeneration and the alignment of our quantification with an independent study clearly supports their identification as RGCs. The selective visibility of RGCs over the abundant displaced amacrine cells suggests that RGCs possess distinct intrinsic optical properties, possibly related to their larger soma size or organelle composition, which render them highly reflective and thus distinguishable with OCT system.

This hypothesis builds on an explanatory framework proposed in previous work to account for the biophysical origins of cellular contrast in OCT. Liu et al. proposed two main factors contributing to RGC soma visibility^15^. The first is a dynamic mechanism rooted in organelle motility, which they described as an intrinsic contrast agent. As organelles move within the cell, the speckle pattern they create changes between successively acquired OCT volumes. Consequently, when dozens of volumes are averaged, this decorrelated intracellular speckle noise is suppressed, while the static, time-invariant outline of the soma is preserved and enhanced. The second factor is a difference in reflectance tied to cellular composition and size. Liu et al. observed that larger somas were systematically brighter, suggesting that their ultrastructure, for instance, a higher density of mitochondria and other organelles characteristic of metabolically active RGCs, contributes to their visibility over smaller, neighboring cells. Taken together, these proposed dynamic and static contrast mechanisms offer a plausible physical basis for the selective visualization of RGCs, distinguishing them from the smaller, less metabolically active, and thus less reflective, displaced amacrine cells within the RGC layer.

While our WSAO OCT platform provides valuable capabilities for in vivo cellular imaging, several limitations should be noted. First, experimental throughput is constrained by time-intensive procedures including AO optimization, volume acquisition (∼100 volumes), and computational processing. Streamlining these steps through automation would enable larger cohort studies. Second, the effective FOV for high-quality AO correction is limited to approximately 300 × 300 µm due to the high curvature of the mouse eye. Future work could incorporate rapid acquisition and mosaicking of multiple AO corrected patches to assess wider retinal regions. Third, our manual approach to RGC quantification is labor-intensive and prone to subjectivity. To address this, we are developing a deep-learning framework for automated cell segmentation. The value of this approach is highlighted by human AO-OCT studies, where automated analysis revealed that RGC soma hypertrophy, a likely indicator of cellular stress and axonal transport failure, combines with density loss to better predict functional decline than conventional metrics^19,20^.

In conclusion, our development of WSAO-OCT overcomes the persistent optical barriers in mouse retinal imaging, enabling, for the first time, visualization of the comprehensive cellular mosaic from the RGC layer to the RPE. This technology opens new avenues for exploring the cellular pathology of retinal diseases in their native context. Ultimately, the ability to repeatedly and non-invasively assess cellular health and degeneration in vivo promises to accelerate the translation of mechanistic insights into therapeutic strategies.

## Materials and Methods

### OCT Imaging System

We employed a custom-developed SD-OCT system (Netra Systems, Inc., Pleasanton, CA, USA) for in vivo mouse retinal imaging (Fig. 6). The system operated across a spectral range of 750-975 nm with a 3 dB bandwidth of 172.4 nm centered at 851.3 nm, delivering 1.4 mW at the cornea. This configuration achieved theoretical resolutions of 1.5 μm axial and 3 μm (1 μm with WSAO) lateral in tissue, with a 1.2 mm collimated beam diameter on the cornea. The system provided 2 mm imaging depth in air with 97 dB sensitivity (Fig. 6). Aberration correction was implemented using a piezoelectric deformable mirror (DM) integrated within a feedback control loop. The DM shape was optimized through sequential modulation of Zernike modes, progressing from lower to higher orders, with an algorithm that maximized image brightness in real-time en face OCT images. An electrically tunable lens (ETL) in the sample arm enabled dynamic focal plane adjustment for selecting different retinal layers of interest (LOI). Image acquisition utilized a high-speed spectrometer equipped with a line-scan camera operating at 100 kHz A-scan rate.

**Figure 6.**
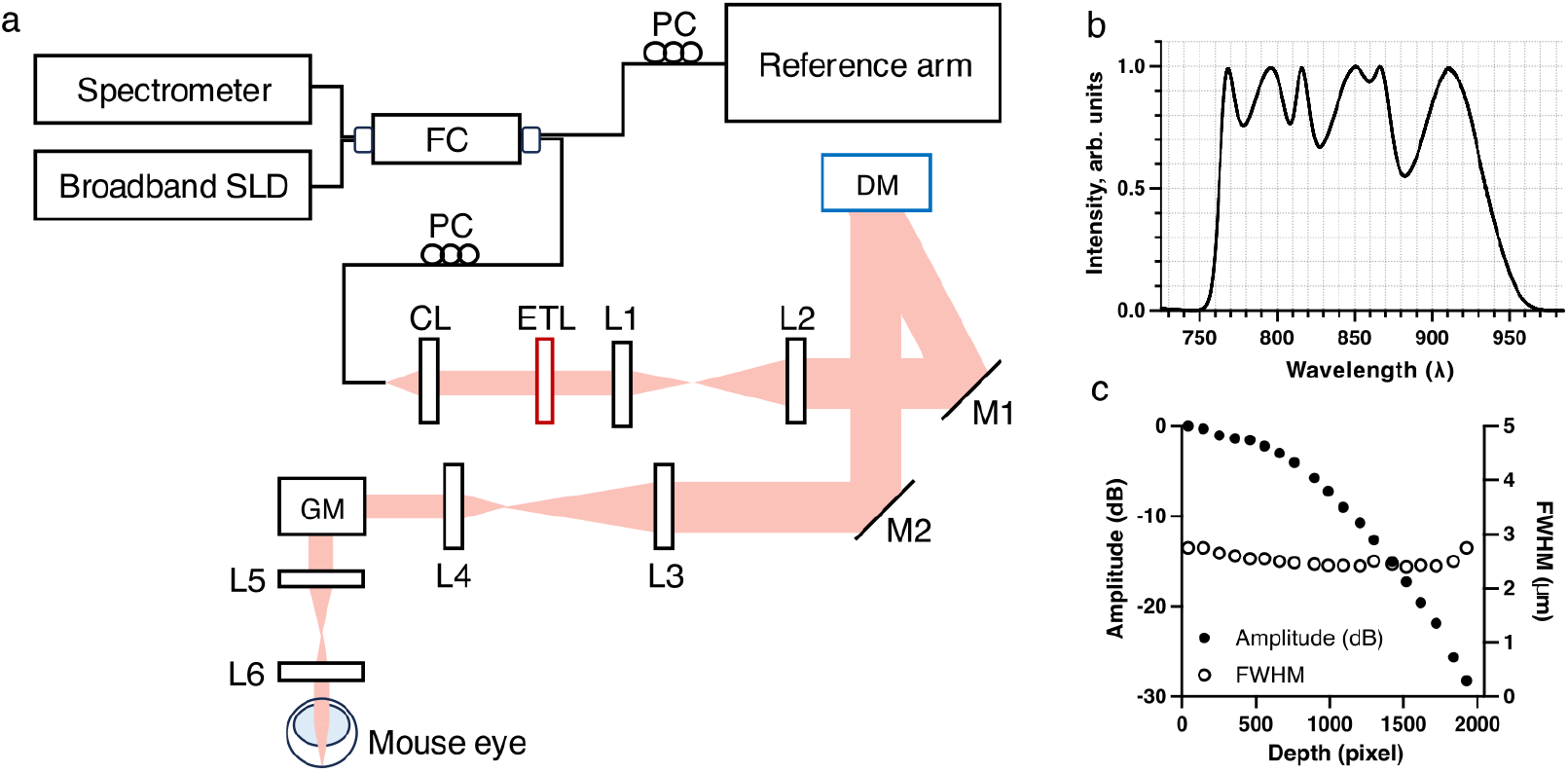
(a) Schematic of the custom-built wavefront sensorless SD-OCT system. SLD: superluminescent diode; FC: fiber coupler; PC: polarization controller; CL: collimation lens; ETL: electrically tunable lens; DM: deformable mirror; GM: galvanometer-scanning mirrors; L1-L6: lenses; M1-M2: mirrors. (b) The emission spectrum of the broadband light source used in the OCT system, indicating the center wavelength (851.3 nm) and bandwidth (750-975 nm). (c) System performance metrics plotted as a function of imaging depth, showing the measured sensitivity roll-off and the axial resolution, defined as the full width at half maximum (FWHM).

### Animal Preparation

All experimental procedures adhered to the Association for Research in Vision and Ophthalmology (ARVO) Statement for the Use of Animals in Ophthalmic and Vision Research and were approved by the Institutional Animal Care and Use Committee (IACUC) at Genentech, Inc. C57BL/6J mice of both sexes, aged 2-4 months, were used for imaging experiments. Anesthesia was induced via intraperitoneal injection of ketamine (100 mg/kg) and xylazine (10 mg/kg). Pupils were dilated with topical 1% tropicamide prior to imaging. To maintain corneal hydration, a custom-designed flanged contact lens was applied with lubricant eye gel^21^. Mice were positioned on a heated platform to maintain body temperature, with head stabilization achieved using a bite bar and flexible tension band to minimize motion artifacts during image acquisition.

### Image Acquisition and Multi-Volume Process

Volumetric OCT data were acquired through raster scanning of retinal ROI. After selecting the ROI and LOI, image quality-based AO correction was applied to compensate for ocular aberrations. Multiple volumes with 512 × 512 A-scans were acquired from identical retinal locations at 2.8 seconds per volume (Fig. 7a). To achieve distinguishable contrast among cell bodies, we developed a multi-stage processing pipeline that enhances signal-to-noise ratio (SNR) through registration, alignment, and averaging of OCT volumes. The workflow processes data in batches of six volumes, chosen to balance processing efficiency with available computational memory, through three sequential stages (Fig. 7b). First, a rigid registration corrects for gross inter-volume misalignments in the B-scan (XZ) plane. For each volume, a 2D side-projection is generated from its central 60 B-scans. These XZ-plane projections are then registered to a common reference to correct for large-scale translational and rotational errors between the volumes. Second, en face registration in the XY plane addresses fine-scale lateral movements. This is achieved by generating en face projections from each volume via standard deviation mapping and then computing and correcting for sub-pixel translations using phase correlation. Third, B-scan alignment corrects for intra-volume axial motion artifacts. This final registration stage analyzes each B-scan independently, calculating and correcting vertical displacements between adjacent frames using column-wise cross-correlation to ensure axial stability. After completing all three registration stages, the aligned volumes within each six-volume batch are averaged to suppress speckle noise while preserving cellular structures. The resulting averaged volumes from each batch are then grouped into new six-volume sets and processed through the same pipeline repeatedly, creating a hierarchical averaging scheme that continues until a single final volume is obtained. Registration accuracy is validated quantitatively through displacement measurements and qualitatively through overlay comparisons at each stage. Detailed implementation is provided in the accompanying pseudocode (Supplementary Fig. S3), and the Python code is available at “https://github.com/unblindness/oct_alignment“.

**Figure 7.**
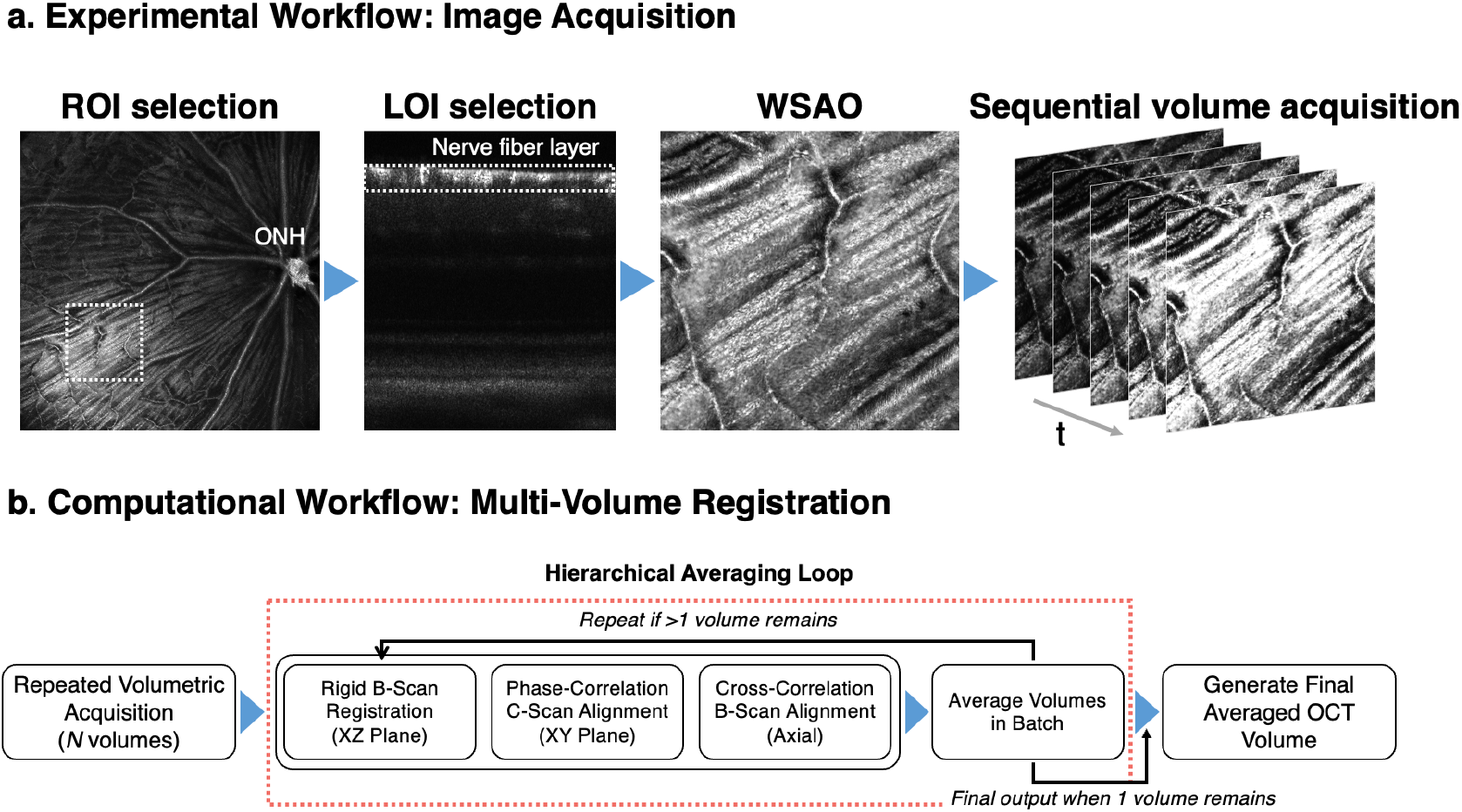
Integrated workflow for cellular-resolution in vivo imaging of the mouse retina. (a) Data acquisition protocol demonstrating the sequential steps from region selection to volume collection, with real-time WSAO optimization applied at each selected retinal layer. (b) Hierarchical computational pipeline that enhances SNR through iterative multi-plane registration (addressing XZ motion, XY drift, and axial displacement) followed by averaging, with the process repeated until convergence to a high-fidelity cellular image.

### Optic Nerve Crush (ONC) Surgery

ONC was performed on the left eye of mice under isoflurane anesthesia (5% induction, 2.5% maintenance, 1.5–3 L/min O2). Following confirmation of surgical anesthesia depth via absent reflexes, mice were positioned on a warming pad. The intraorbital optic nerve was accessed through conjunctival incision and blunt dissection of muscles and connective tissue. Using self-closing jeweler’s forceps (Dumont #5, Fine Science Tools, Foster City, CA, USA), the nerve was crushed 1 mm posterior to the globe for 10 seconds, carefully preserving meningeal sheaths and blood supply. After repositioning the eye, ophthalmic antibiotic ointment was applied to the surgical site and lubricant ointment to the contralateral eye. Mice were monitored until fully recovered and observed daily thereafter for any adverse effects.

### Immunohistochemistry of Whole-Mount Retina

Mice were euthanized by CO_2_ inhalation followed by cervical dislocation. Eyes were enucleated and fixed in 4% paraformaldehyde for 2 hours at room temperature. After dissection, retinas were washed in PBS and blocked for 2 hours in staining buffer (10% normal donkey serum, 0030-01, Southern Biotech, Birmingham, AL, USA; 0.5% Triton X-100, Sigma-Aldrich, St. Louis, MO, USA in PBS). Retinas were incubated with rabbit anti-RBPMS primary antibody (1:200, NBP2-20112, Novus Biologicals, Littleton, CO, USA) for 48 hours at 4°C. Following three PBS washes, retinas were incubated for 48 hours at 4°C with a cocktail containing goat anti-rabbit IgG FITC (1:100, 31635, Thermo Fisher Scientific, Waltham, MA, USA), NeuroTrace™ 435/455 Blue Fluorescent Nissl Stain (1:100, N21479, Thermo Fisher Scientific, Waltham, MA, USA), and Alexa Fluor^®^ 647 anti-mouse CD31 antibody (1:100, 102516, BioLegend, San Diego, CA, USA). After final PBS washes, retinas were flattened with four radial cuts, mounted using ProLong™ Glass Antifade Mountant (P36982, Thermo Fisher Scientific, Waltham, MA, USA), and imaged using a Leica SP8 confocal microscope.

### Image Quality Assessment

The effectiveness of AO correction was quantified using two complementary metrics applied to en face OCT images: Tenengrad (edge sharpness) and intensity standard deviation (local contrast), where higher values indicate better image quality. For each 20×20 pixel patch of a grayscale image, the Tenengrad metric *T*(*P*) = ∑_*i,j*_|∇*I*(*i, j*)| quantified local sharpness, where |∇*I*(*i, j*)| represents the gradient magnitude at pixel position (*i, j*) derived from Sobel operators. Local contrast was assessed via intensity standard deviation 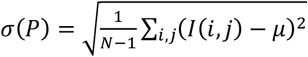, where *I*(*i, j*) represents pixel intensity, *µ* the patch mean intensity, and *N* the total number of pixels in the patch. These metrics were computed across non-overlapping patches to generate spatial quality maps. Computation for these two metrics was performed in MATLAB R2020a (MathWorks, Natick, MA, USA).

The effectiveness of volume averaging for speckle noise suppression was quantified using three distinct metrics. First, normalized speckle contrast was calculated for each averaged volume to directly measure noise reduction. Speckle contrast was defined as the ratio of standard deviation (σ) to mean intensity (μ) within the ROI: C = σ/μ. Values were normalized to the single-volume image (X = 1) as reference (normalized value = 1.0), enabling direct comparison of noise reduction across different averaging levels. SSIM was computed to assess the convergence of image quality relative to the fully averaged dataset.

Using the 200-volume averaged image as the reference standard, SSIM values were calculated for each volume average (X = 1 to 200). SSIM values range from -1 to 1, with 1 indicating perfect structural similarity to the reference image. In addition, line intensity profiles were extracted from en face OCT images. A linear line of interest was manually selected across multiple representative cell bodies in the reference image (200-volume average). The same line coordinates were applied to all images in the series to ensure consistent sampling. Cross-correlation analysis was performed to assess similarity between line profiles from different volume averages. Using the 200-volume averaged image as the reference standard, Pearson correlation coefficients were calculated between each test profile and the reference profile. One-phase exponential decay models were fitted to all metrics to characterize the relationship between volume averaging and image quality, determining the optimal number of volumes for practical imaging applications. The analyses, including the calculation of speckle contrast, SSIM, and Pearson correlation coefficients, were performed using Python (v3.9) with scientific computing libraries such as NumPy, SciPy, and Scikit-image.

### Manual Cell Counting

Manual cell counting was performed on three sets of images: in vivo OCT en face volumes of the RGC layer, and ex vivo whole-mount retinas stained for RBPMS and NeuroTrace. For all image types, analysis was conducted using ImageJ (National Institutes of Health, USA). A grid overlay was applied to each field of view to divide the image into smaller, uniform sub-regions, enabling systematic and unbiased counting within each sub-region. For OCT volumes, each sub-region was manually scanned through the z-stack. A structure was identified and counted as a cell only if it exhibited a round, soma-like morphology and was visible across a minimum of three consecutive stacks. For the RBPMS and NeuroTrace fluorescence images, a cell was counted if it displayed a clear cellular shape and was positively labeled with the corresponding fluorescent marker.

### GCC Thickness Measurement

GCC thickness was quantified using a custom Python-based semi-automated segmentation algorithm. Following preprocessing with image enhancement and vertical derivative filtering, the user identifies a single A-line in the central B-scan and marks two features: an intensity peak (anterior RNFL) and valley (posterior inner plexiform layer (IPL) boundary). The algorithm then searches within ±2-pixel axial windows to detect the strongest local extrema column-by-column, generating smooth boundary curves across the initial B-scan. These seed curves propagate through adjacent frames via a surface-growing approach, where each previous boundary guides the search in subsequent slices. The resulting 3D segmentation yields voxel-wise GCC thickness maps computed as the axial distance between RNFL and IPL boundaries.

### Statistical Analysis

All statistical analyses were performed in GraphPad Prism 9 (GraphPad Software, San Diego, CA, USA) and are reported as mean ± standard deviation (SD). Longitudinal changes in cell counts and GCC thickness measured from OCT images at five matched time points (baseline; 1, 3, 5 and 7 days post-ONC) were assessed with repeated-measures one-way ANOVA using the Geisser–Greenhouse correction for degrees of freedom. When the overall F-test reached significance, Dunnett-adjusted post-hoc tests compared each post-ONC time point with its respective baseline while controlling the family-wise error rate. Two-tailed, multiplicity-adjusted P ≤ 0.05 was regarded as statistically significant.

## Data availability

The data supporting the findings of this study are available from the corresponding author upon reasonable request.

## Code availability

Code for multi-volume registration and averaging is available at [https://github.com/unblindness/oct_alignment].

## Author Contributions

T-H.K. designed the study, developed the volume registration algorithm, performed the experiments, analyzed the data, and wrote the original manuscript draft. R.W. contributed to the study’s conceptual development and manuscript revision. J.E. supervised the project, provided technical guidance, and contributed to manuscript revision.

## Conflict of Interest

T-H.K., R.W., and J.E. are employees of Genentech, Inc. The authors declare no other conflicts of interest.

## Notes

https://github.com/unblindness/oct_alignment

