## Supplemental Information for "Cellular-resolution OCT reveals layer-specific retinal mosaics and ganglion cell degeneration in mouse retina in vivo"

**This PDF file includes:**
Supplementary Figures S1 to S3.
Legends for Movie S1.

**Other supplementary materials for this manuscript include the following:**
Movie S1.mpeg

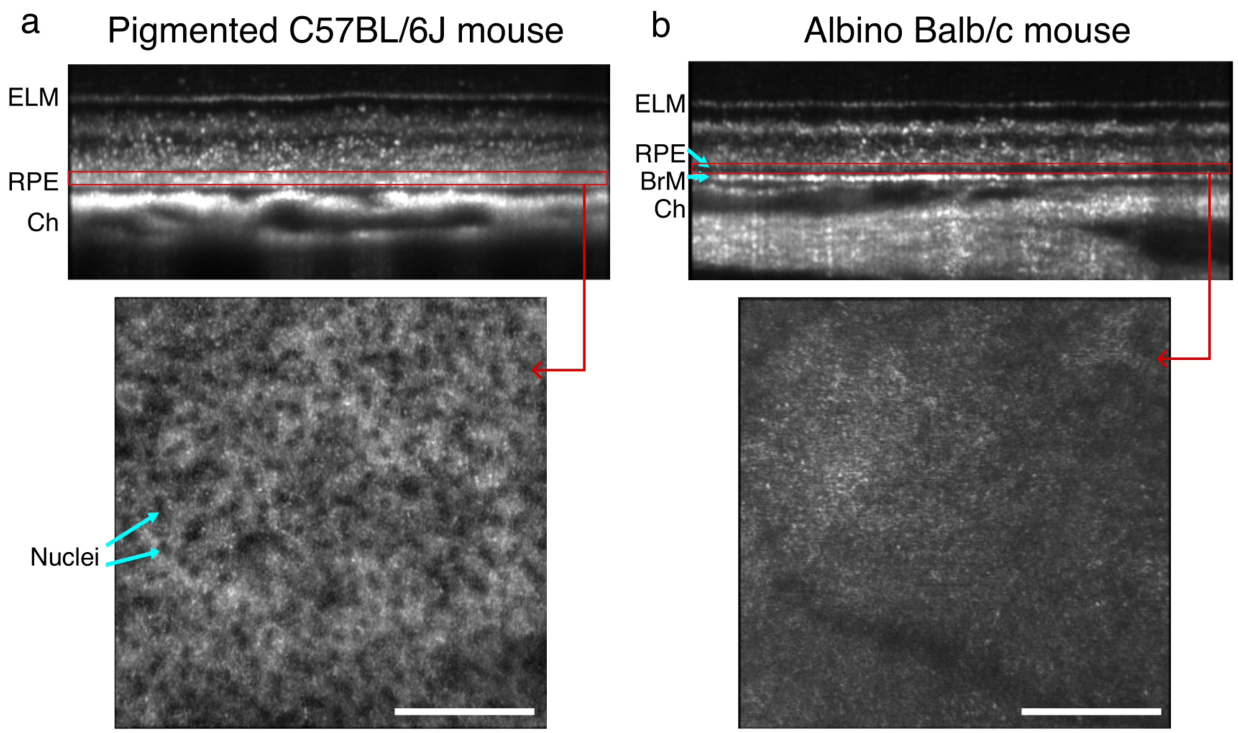


**Supplementary Figure S1.** The RPE mosaic pattern is visible only in pigmented mice. Representative cross-sectional (top) and *en face* (bottom) OCT images of the RPE layer in (a) a pigmented C57BL/6J mouse and (b) an albino Balb/c mouse. In the pigmented mouse, the *en face* image reveals a distinct mosaic pattern of regularly spaced, hyporeflective spots, presumed to be nuclei enveloped in hyporeflective melanosomes, surrounded by hyperreflective cytoplasm. This cellular mosaic is absent in the non-pigmented RPE of the albino mouse. (ELM, external limiting membrane; RPE, retinal pigment epithelium; BrM, Bruch's membrane; Ch, Choroid). Scale bars: 100 µm.

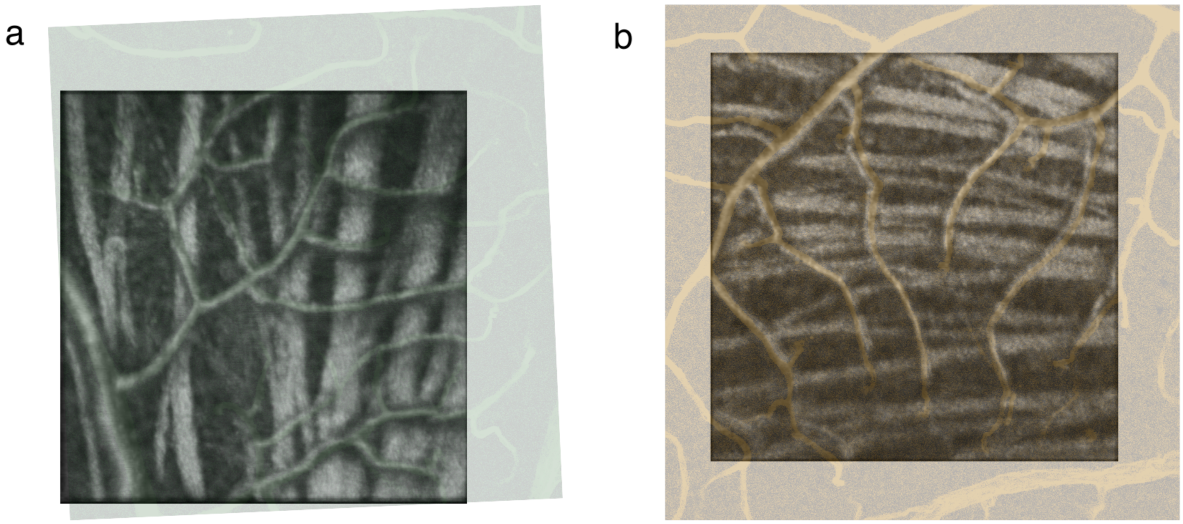


**Supplementary Figure S2.** Calibration of the OCT field of view (FOV) using co-registered confocal images of retinal vasculature. To determine the precise FOV of the in vivo OCT imaging, a whole-mount retina was stained for vasculature with a CD31 marker and imaged via confocal microscopy. The resulting confocal image, which has a known physical size of 388 × 388 µm, was then overlaid onto the corresponding OCT image using the vascular pattern for alignment. The overlay is shown for two independent samples using different pseudocolors for visualization purposes: green in (a) and yellow in (b). By using the known dimensions of the confocal image as a calibrated ruler, the FOV of the underlying OCT image was estimated to be approximately 310 × 310 µm, which corresponds to roughly 8 x 8 degrees of the adult mouse visual field.

**Input:** A set of $N$ initial OCT volume files
**Output:** A single, final aligned and averaged volume

Let $V_{current}$ be the initial set of $N$ volume files
**while** number of volumes in $V_{current} > 1$ **do**
    Let $V_{next\_level}$ be an empty list
    **for** each batch of volumes in $V_{current}$ **do**
        **Load and preprocess** volumes
    *Stage 1: Rigid Registration (XZ Plane)*
        **Compute** side-projection from central 60 B-scans
        **Perform** rigid registration of side-projections
        **Apply** transformations to original volumes
    *Stage 2: En Face Registration (XY Plane)*
        **Generate** En Face projections and apply translational alignment
    *Stage 3: B-scan Alignment (Axial)*
        **Concatenate** volumes into an interleaved volume
        **Apply** vertical displacement correction to B-scans
    *Averaging and Saving for the Batch*
        **Average** aligned frames to produce $V_{averaged}$
        **Save** the averaged volume $V_{averaged}$
        **Add** $V_{averaged}$ to list $V_{next\_level}$
    **end for**
    $V_{current} \leftarrow V_{next\_level}$ ▷ Set outputs as inputs for next level
**end while**
**return** the single volume remaining in $V_{current}$

**Supplementary Figure S3.** Hierarchical multi-volume registration and averaging algorithm. The iterative pipeline processes OCT volumes through three-stage registration (rigid XZ-plane, en face XY-plane, and axial B-scan alignment) followed by averaging to progressively reduce volume count while enhancing image quality through speckle noise suppression.

**Supplementary Movie S1.** In vivo visualization of cellular mosaics in four distinct retinal layers. This supplementary file consists of four separate movies, each presenting an axial fly-through within a specific retinal layer of a living mouse. The movies display the distinct cellular mosaics of the retinal ganglion cell (RGC) layer, the inner nuclear layer (INL), the outer nuclear layer (ONL), and the retinal pigment epithelium (RPE), respectively. Together, these visualizations demonstrate the capability of the integrated workflow to resolve individual cells with their unique layer-specific morphologies in vivo. Scale bar, 100 µm.
